# Decreased BOLD signal variability in middle-aged and older adults on the Autism Spectrum

**DOI:** 10.1101/2024.12.21.629918

**Authors:** Stephanie Pedrahita, Annika Linke, Michaela Cordova, Molly Wilkinson, Janice Hau, Gioia Toro, Kalekirstos Alemu, Jiwandeep Kohli, Ralph-Axel Müller, Ruth Carper

## Abstract

**Purpose:** Autism spectrum disorder (ASD) is a lifelong neurodevelopmental disorder. Preliminary evidence suggests an increased risk for accelerated or early-onset cognitive and neurological decline in ASD. While it is well established that brain development in children, adolescents and young adults with ASD diverges from neurotypical (NT) peers, it is unknown how brain function is impacted in older adults with ASD. Understanding age-related changes of brain function in ASD is crucial to establish best practices for cognitive and health screenings in adults with ASD and develop interventions that might reduce the risk of accelerated decline. Decreases in blood-oxygenation-level-dependent (BOLD) signal variability (BSV) in typical aging have been shown across multiple studies, likely reflecting declining Gamma-Aminobutyric Acid (GABA) activity, and is associated with poorer cognitive performance. We hypothesized that adults with ASD would show reduced BSV compared to the NT group, with steeper negative age associations in the ASD than NT group.

**Methods:** The study assessed BSV in a cohort of adults (40-70 years), 28 with ASD and 39 age-matched NT. General linear models tested for main effects of diagnostic group (ASD, NT), age and group-by-age interactions (controlling for RMSD).

**Limitations:** Our cross-sectional data and small sample size highlight the need for longitudinal analyses in larger cohorts, alongside exploring links to cognitive function. Additionally, psychotropic medications used by our cohort of adults on the autism spectrum may have affected BSV.

**Results:** Significant group-by-age interactions were observed for the right insular, left temporal occipital fusiform, right frontal orbital and right inferior lateral occipital cortex, with BSV showing strong negative associations with age in the ASD but not NT group.

**Conclusion:** These findings suggest that BSV decreases may occur earlier in adults on the autism spectrum compared to their neurotypical peers, possibly indicating accelerated aging. However, given limited prior research, additional longitudinal analyses will be necessary to determine if the results presented truly reflect accelerated aging or arise from lifelong persistent differences in brain function.

## Background

Autism spectrum disorder (ASD) refers to a lifelong neurodevelopmental disorder characterized by— (1) impairments in social communication or interaction across multiple contexts and (2) restricted, repetitive patterns of behavior, interests, or activities (American Psychiatric Association, 2013). These characteristics are independent of culture, race, ethnicity, or socioeconomic group (Lord et al., 2012). Most ASD research and particularly neuroimaging research has been focused on better understanding ASD in early development, but these symptoms and behaviors have been shown to persist into adulthood. Multiple studies have also identified differences in cortical morphology, diffusion, MEG and functional MRI in children and adults with ASD compared to their neurotypical (NT) peers across the lifespan (Gu et al., 2024; Hau et al., 2022; Wilkinson et al., 2022; Van Rooij et al., 2018). While it is well established that brain development in children, adolescents and young adults with ASD diverges from NT peers, far less is known about how brain function is impacted in older adults with ASD and what consequences this may have for cognition and behavioral abilities. Given preliminary evidence suggesting an increased risk for accelerated or early-onset cognitive and neurological decline (Mason et al., 2021; Torres et al., 2020; Walsh et al., 2022), it is important to better understand the relationship between brain function and behavior in autism in late adulthood.

One measure of brain function that has been associated with typical aging is blood-oxygenation-level-dependent (BOLD) signal variability (BSV), measured by functional magnetic resonance imaging (fMRI) through the blood oxygen-level dependent (BOLD) signal. While once considered “noise” in fMRI, BSV has been shown to change with age and vary across different cognitive states (Garrett et al. 2010, 2011). Previous studies suggest that higher BSV may allow for a wider range of responses, making the brain more flexible (Grady & Garrett, 2018, McDonnell & Ward, 2011; Burzynska et al., 2015). This broader range of responses might facilitate the brain’s ability to process and respond to a wider variety of environmental inputs. It has also been found that BSV is lower in areas that affect memory performance, cognition, and dynamic functioning of the brain (Garrett et al., 2014; Guitart-Masip et al., 2016; Protzner et al., 2013).

BSV decreases in typical aging, (Garrett et al., 2011; Grady & Garrett, 2018; Waschke et al., 2021). Specifically, Garret et al. (2010) found that at rest, older brains showed less BSV than younger brains across a broad subset of regions. In a separate study, they found that healthy younger adults (20–30 years) showed higher BOLD variability across cognitive tasks and greater variability-based regional differentiation compared with older, poorer-performing adults (56–85 years) (Garret et al., 2011). In a large cohort of neurotypical children and adults, Nomi et al. (2017) found BSV decreased from ages 6 to 85 years in the majority of brain regions examined, suggesting age-related differences in brain flexibility that may underlie behavioral and cognitive changes with age.

To better understand the mechanism underlying age-related declines in BSV, Lalwani et al. (2021) tested the hypothesis that changes in GABA activity are driving the observed reduction in BSV with age. They were motivated by previous pharmacological findings indicating that reduced GABA activity in animal models decreases network signal variability and the number of states the cortical network can visit, as well as previous work showing that older adults show lower GABA levels and lower BSV across cortex (Chamberlain et al., 2019; Cassady et al., 2019; Gao et al., 2013; Garrett et al., 2011, 2014; Grady & Garrett, 2018; Porges et al., 2017; Lalwani et al., 2019; Shew et al., 2011; Waschke et al., 2021). Lalwani et al. (2021) quantified BSV in 20 young (age 18–25 years) and 24 older (age 65–85 years) healthy human adults during placebo and GABA agonist sessions and again found that older adults exhibited lower signal variability at placebo. GABA-related boosts in BSV were largest in the poorest performing older adults. This suggests that GABA plays a role in BSV with consequences for cognition in older adults.

In children on the autism spectrum, GABA activity has been found to be reduced by several groups (Gaetz et al., 2014; Harada et al, 2011; Puts et al., 2017; Kubas et al., 2012). A number of the genetic differences associated with ASD affect GABA function by impacting interneuron numbers, GABA synthesis or GABA receptors (Lee et al., 2017), leading to imbalances in excitation and inhibition commonly observed in animal models of the disorder.. GABA decreases in typical aging, as does BSV. It is therefore important to understand how the combined effects of aging and autism affect BSV, and how this relationship impacts cognitive function in older adults on the autism spectrum. To our knowledge, only one study has examined BSV in autism, including 20 boys on the autism spectrum (M=13.25 years) and 17 neurotypical boys (M=13.42 years) (Easson & McIntosh, 2019). They found that greater severity of ASD-like behaviors was associated with decreased BSV. In the present study, we hypothesized that older adults with ASD would show reduced BSV compared to the NT group, with steeper negative age associations in the ASD than NT group.

## Methods

### Participants

This study assessed BSV cross-sectionally in a cohort of adults (40-70 years), 28 with ASD and 39 NT, who participated in a multimodal longitudinal study of aging in ASD (Table 1). There were no significant differences between the ASD and NT groups on age, gender, non-verbal IQ, body mass index, or co-occurring hypertension. A list of medications for all participants can be found in Supplementary Table S5. There was a significant difference found for head motion (RMSD), which was added as a covariate to all analyses. ASD diagnoses were made by a licensed clinical psychologist in accordance with the DSM-5 (American Psychiatric Association, 2013) criteria and supported by Module 4 of the Autism Diagnostic Observation Schedule, Second Edition (ADOS-2) (Lord et al., 2012) and a semi-structured clinical interview to probe for developmental history and ASD-relevant difficulties during school-age. For both ASD and NT groups, participants with a known history of neurologic (e.g., epilepsy, tuberous sclerosis) or genetic (e.g., fragile X, Rett syndrome) conditions other than autism were excluded. NT participants were excluded for personal or family history of ASD or personal history of other neurologic conditions or serious mental illness.

**Table 1.** Demographic information for NT and ASD groups.

|  | NT (N= 39) |  | ASD (N= 29) |  | Statistics |
| --- | --- | --- | --- | --- | --- |
| Gender | 7 females | | 7 females | | $\chi^2=0.5$, $p=0.5$ |
|  | **mean ± SD** | **range** | **mean ± SD** | **range** |  |
| Age (y) | 52.4±7.5 | 40.3-69.9 | 51.5±7 | 40.3-67.4 | t(66)=0.6, $p=0.6$ |
| RMSD | 0.1±0.04 | 0.04-0.2 | 0.1±0.05 | 0.04-0.2 | t(66)=-2.8, $p<0.006^*$ |
| BMI | 26.7±3.5 | 22.5-34.2 | 29.7±5.1 | 22.9-41.2 | t(57)=-2.7, $p=0.01^*$ |
| WASI VCI | 113±13.4 | 85-144 | 101.2±23.7 | 45-160 | t(66)=2.5, $p<0.05^*$ |
| WASI PRI | 109.2±12.3 | 75-138 | 106.8±21.1 | 63-141 | t(66)=0.6, $p=0.56$ |
| WASI FSIQ | 112.7±11.8 | 90-138 | 104.3±22.4 | 58-143 | t(66)=2, $p=0.05^*$ |
| Mean HR | 63.4±10.3 | 46.3-84 | 67.1±10.8 | 38-88.5 | t(66)=-1.4, $p=0.2$ |
| Mean STD HR | 1.9±0.9 | 0.8-5.3 | 2.2±1.1 | 0.6-4.6 | t(64)=-0.9, $p=0.3$ |
| Mean RVT | 0.5±0.2 | 0.2-0.8 | 0.4±0.2 | 0.1-0.8 | t(63)=1.4, $p=0.2$ |
| Mean STD RVT | 0.1±0.04 | 0.06-0.2 | 0.1-0.05 | 0.07-0.3 | t(63)=-1.2, $p=0.2$ |

Abbreviations: RMSD=Root-Mean-Square Deviation; BMI=Body Mass Index; WASI=Wechsler Abbreviated Scales of Intelligence; VCI= Verbal Comprehension Index; PRI=Perceptual Reasoning Index; HR=Heart Rate; RVT= Respiratory Volume Per Time. Data were missing for several participants: Mean HR (1 NT, 1 ASD), Mean STD HR (1 NT, 1 ASD), Mean RVT (1 ASD, 2 NT), Mean STD RVT (1 ASD, 2 NT), BMI (3 NT, 6 ASD).

### Data Acquisition

Participants completed an MRI session at the University of California San Diego Center for Functional MRI on a 3T GE Discovery MR750 scanner using a Nova Medical 32-channel head coil. A multiband EPI sequence allowing for simultaneous acquisition of multiple slices was used to acquire two 6 minute fMRI runs with high spatial and temporal resolution (TR=800ms, TE=35ms, flip angle 52°, 72 slices, multiband acceleration factor 8, 2 mm isotropic voxel size, 104×104 matrix size, FOV 20.8cm, 400 volumes per run). Two separate 20s spin-echo EPI sequences with phase encoding in both A>>P and P>>A directions were acquired using the same matrix size, FOV and prescription to correct for susceptibility-induced distortions. Prior to their functional scans, participants received the instruction: “Keep your eyes on the cross. Let your mind wander, relax, but please stay as still as you can. Try not to fall asleep.” Adherence to the instructions to remain awake with eyes open was monitored with an MR-compatible video camera and only participants compliant with these instructions were included in analyses. Heart rate and respiration were recorded during the functional scans using a Biopac pulse oximeter placed on the right index finger and respiration was measured using a breathing belt placed across the participant’s diaphragm. Physiological recordings during the fMRI scans were analyzed with the PhysIO toolbox (Kasper et al., 2017). The mean and standard deviation of heart rate and respiratory volume were calculated for each participant. Groups did not differ on any of the physiological measures except for BMI (see Table 1). Additional regression analyses between BSV and BMI were performed with no significant correlations (*p*<.05) identified. Physiological measures were therefore not included as covariates in subsequent analyses.

### fMRI preprocessing and denoising

fMRI data underwent standard preprocessing, denoising, and were analyzed in Matlab 2016b (Mathworks Inc., Natick, MA, USA) using SPM12 (Wellcome Trust Centre for Neuroimaging, University College London, UK) and the CONN toolbox v17f., including rigid-body realignment, normalization to the MNI template, bandpass filtering, and nuisance regression to remove physiological and motion confounds. Functional images were then corrected for susceptibility-induced distortions using the two spin-echo EPI acquisitions with opposite phase encoding directions and FSL’s TOPUP tools (Smith et al., 2004). Then, functional images were motion-corrected using rigid-body realignment through SPM12. The Artifact Detection Toolbox (ADT, as installed with CONN v17f) identified outliers in the functional image time series from the resulting 6 motion parameters (3 translational and 3 rotational) that had frame-wise displacement (FD) >0.9mm and/or changes in signal intensity that were greater than five standard deviations. Since oscillations due to respiration are prominent in motion parameters derived from multiband EPI realignment (Fair et al., 2020) and would result in unnecessary censoring of large chunks of data in some participants, the thresholds to detect outliers were more lenient than those used for standard resting state fMRI acquisitions with slower TRs.

### Quantification of brain signal variability

Since all analyses were run on averaged voxel time series within pre-defined ROI, no prior smoothing was applied to the data. Voxel time series were normalized to percent-signal change in each run. Band-pass filtering using a temporal filter of 0.008 to 0.08 Hz was carried out as part of the nuisance regression (“simult” option in the conn toolbox) which also included scrubbing of the motion outliers detected by the ART toolbox, and regression of the 6 motion parameters and their derivatives, as well as the first five Principal Component Analysis (PCA) component time series derived from the CSF and white matter masks. The residuals of the nuisance regression were then used for all subsequent resting state analyses with timeseries concatenated across the two runs. Average BSV series were extracted from the Harvard-Oxford anatomical parcellation and BSV calculated as the standard deviation of the timeseries for each ROI as in Lalwani et al. (2021). ROIs were those identified by Lalwani et al., (2021) to show significant age-related reductions in BSV and included 14 frontal, 12 temporal, 2 parietal, 6 occipital and 2 insular cortical areas. See Supplementary Tables S1-2 for all ROIs tested.

### Statistical analyses

General linear models tested for main effects of diagnostic group (ASD, NT), age and group-by-age interactions (controlling for RMSD) in each ROI. Results are reported at a *p*<0.05 uncorrected *p*-value threshold. Multiple-comparison corrected statistical significance was defined as Benjamini-Hochberg FDR-adjusted *p*<0.1.

## Results

For the main effect of age, all ROIs showed a decrease in BSV (Figure 1) which was significant for 12 of the ROIs at an uncorrected p-value threshold of *p*<.05. After applying the Benjamini-Hochberg false discovery rate (BHFDR) correction, we found that the differences remained statistically significant for three regions: right insular cortex, left temporal occipital fusiform cortex, and right frontal orbital cortex. Detailed statistics for the main effect of age for all ROIs can be found in Supplementary Table S1.

**Figure 1.**
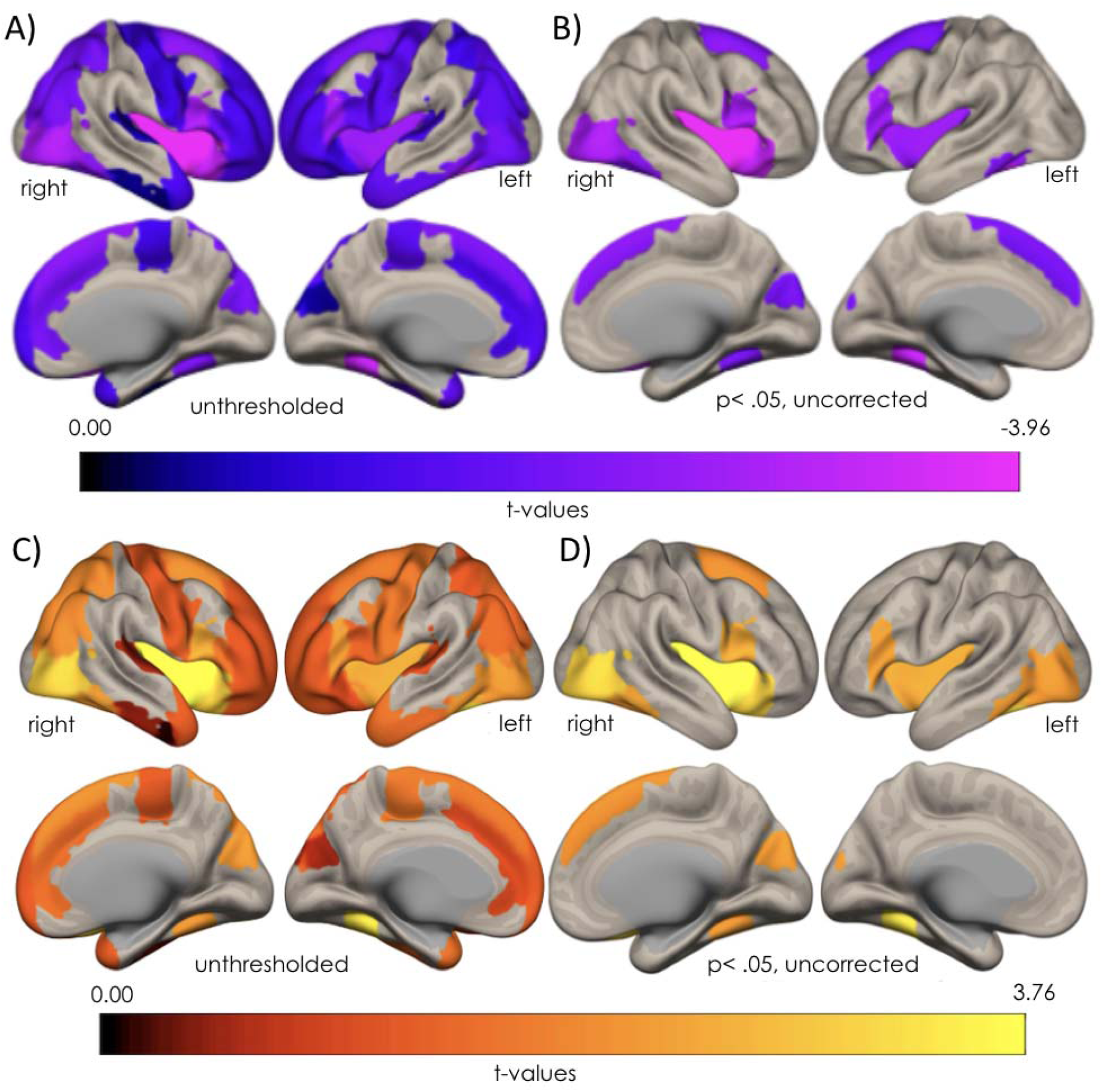
BSV decreased with age across groups for all included ROIs: (A) *t*-values for main effect of age for all ROIs included in analyses, and (B) 4 ROIs (right insular cortex, left temporal occipital fusiform cortex, right frontal orbital cortex and right lateral occipital cortex, inferior division) showing an effect at *p*<.0.5 (uncorrected). (C) Age-by-Diagnosis interaction results (*t*-values) for all ROIs included in analyses, and (D) ROIs that were significant at a *p*-value of *p*<.0.5 (uncorrected).

For the main effect of diagnosis, 12 ROIs showed a significant effect at a *p*<.05 uncorrected p-value threshold. After applying BHFDR correction to account for multiple comparisons, we found that the differences remained statistically significant for four regions: right insular cortex, left temporal occipital fusiform cortex, right frontal orbital cortex and right lateral occipital cortex, inferior division, with older adults on the autism spectrum demonstrating lower BSV compared to their neurotypical peers. See Supplementary Table S2 for statistics for all ROIs tested.

For the interaction between age and diagnosis, 13 ROIs showed effects a *p*<.05 (uncorrected; Figure 1). For all these interactions, ROIs showed a decrease in BSV for the ASD but not NT group (Figure 2). After applying the BHFDR correction to account for multiple comparisons, we found that the differences remained statistically significant for four regions: right insular cortex, left temporal occipital fusiform cortex, right frontal orbital cortex and right lateral occipital cortex, inferior division. For the ASD group, all of these regions showed a statistically significant negative correlation with age; for the NT group only the left temporal occipital fusiform cortex and the right lateral occipital cortex, inferior division showed significant correlations with age, and these were in the positive direction (Figure 2). See Supplementary Table S3 for statistics for all ROIs tested.

**Figure 2.**
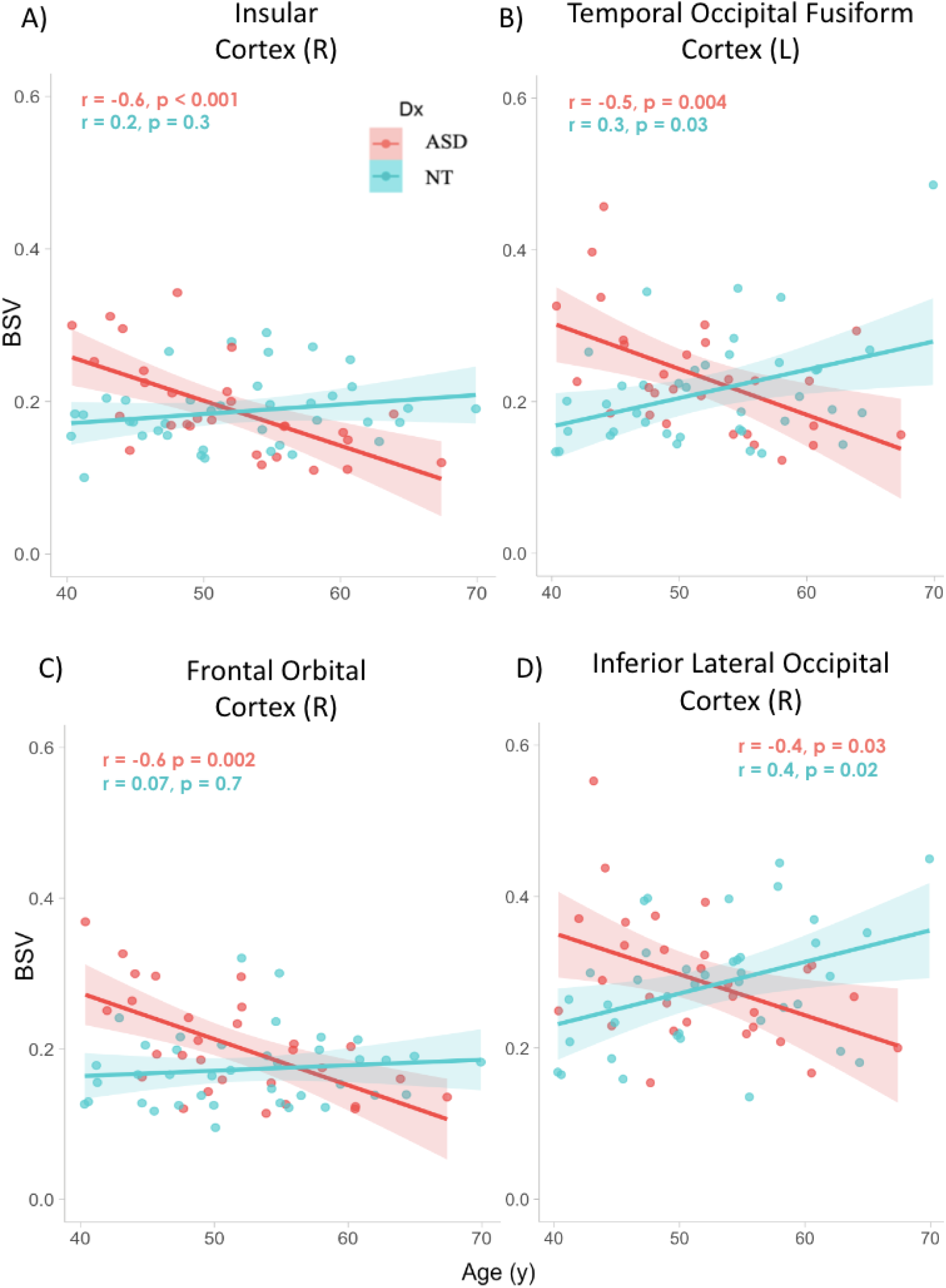
Age-by-Group interactions for the four ROIs that remained significant after BHFDR correction for multiple comparisons in A) right insular cortex, B) left temporal occipital fusiform cortex, C) right frontal orbital cortex, and D) right lateral occipital cortex, inferior division. BSV declined with age in the ASD but not NT group in all ROIs.

## Discussion

In the present study, BSV was investigated in middle-aged and older adults with ASD and compared to their neurotypical peers. We observed that BSV showed negative correlations with age across all included ROIs (with significant decreases with age for 12 regions), replicating previous findings in typical aging (Lalwani et al. 2021; Kielar et al., 2016; Nomi et al., 2017; Grady & Garrett, 2018). The regions showing the strongest age-related effects (remaining significant after adjustment for multiple comparisons) were the right insula, left temporal occipital fusiform cortex, right frontal orbital cortex, and lateral occipital cortex inferior division. Additionally, our finding of steeper age-related BSV decrease in adults with ASD compared to NT adults in several ROIs suggests accelerated brain aging in ASD. While these results were cross-sectional and may thus reflect persistence of earlier differences in brain development, other researchers have found no group differences in BSV between children and adolescents with ASD when compared to age-matched NT peers (Easson & McIntosh, 2019). This supports the interpretation that our findings of decreasing BSV with age in ASD adults might reflect accelerated or earlier-onset brain aging.

While the age-related effects and age-by-group interactions were consistent across all ROIs, the regions with the strongest effects (surviving multiple comparison correction) included areas of the brain frequently implicated in both aging and ASD. For example, multimodal neuroimaging has repeatedly implicated the insula in both aging neurotypical adults and in younger individuals on the autism spectrum (Kleinhans et al., 2016; Nomi et al., 2019; Reiter et al., 2019; Wallace et al., 2010). The insular cortex is thought to modulate social communication skills, including perceiving, imitating, and empathizing with others’ emotional expressions, as well as speech and language functions (Carr et al., 2003; Oh et al., 2014). Researchers have found atypical right insular function in ASD during socio-emotional processing tasks compared to their neurotypical peers for both children and adult populations (Barttfeld et al., 2012; Odriozola et al., 2016). Additionally, ASD participants with higher sensory and communication support needs exhibit overconnectivity in the right insula compared to ASD participants with lower sensory and communication support needs and their neurotypical peers (Reiter et al., 2019).

The main effects of age observed in the current study seem to be driven by the ASD participants (i.e. BSV did not decrease in the NT group). While this seems to contradict previous reports by other researchers who have found significant age-related declines of BSV in NT populations (Lalwani et al. 2021, Nomi et al., 2017), we suspect that this is due to the comparatively young age of participants in our study (mean 52 years). For example, Nomi et al., (2017) observed the steepest declines in their quadratic model in participants older than 60 years. Similarly, the older adult group studied in Lalwani et al. (2021) only included adults aged ≥65 years. The finding that decreases with BSV are already observed in middle-aged autistic adults, thus, further points towards potentially accelerated or earlier onset brain aging in ASD.

It is important to better understand what might be causing lower BSV, and a steeper age-related BSV decrease in middle- and older-age adults with ASD. This relationship could be related to changes in GABA, as discussed by Lalwani et al. (2021), or be due to other physiological differences that exist between ASD and NT groups. For instance, Tsvetanov et al. (2021) found that, when controlling for both cardiovascular and cerebrovascular effects, an age effect was no longer seen for resting-state BSV. Therefore, joint consideration of these measures is necessary for understanding the age effect differences found between ASD and NT participants. Researchers have also found that physiological processes that change in aging (e.g. heart and respiratory rate) can impact cerebral blood flow as well as pulsatory artifacts (Birn et al., 2006; Chang et al., 2009; Glover & Ress, 2000; Song et al., 2023; Tsvetanov et al., 2021).

These factors, in turn, influence BSV, and might confound age-related effects (Grady & Garrett 2014; Kannurpatti et al., 2010; Tsvetanov et al., 2021). Although cardiovascular and cerebral vascular health risks may play an important role in the relationship between BSV and aging, other researchers also propose that underlying mechanisms such as inhibitory neurotransmitters (e.g. GABA) can be a driving force for the interaction between BSV and aging (Lalwani et al., 2021).

The findings of the present study are unlikely to be driven by differences in physiological variables (such as respiration) since there were no significant differences between our groups on these measures. However, we cannot rule out the possibility that co-occurring medical conditions, such as high blood pressure, which may be more prevalent in older adults with ASD, might partially affect the results. Lower BSV observed in adults with ASD could also be due to differences in cardiovascular health. Previous researchers have found that older adults with ASD are at a higher risk of cardiovascular disease compared to their NT peers (Dhanasekara et al., 2023). Supplementary analyses show no group differences for relevant comorbidities for BSV (Supplementary Table S4), however, information on health status was incomplete for a portion of subjects (Supplementary Table S4, Note). Importantly, if effects were driven by increased cardiovascular comorbidity in ASD, this could serve as a point for intervention through pharmacological and behavioral means, which deserves future study.

Psychotropic medications commonly taken by older adults on the autism spectrum may also influence BSV, which is a typical limitation in most neuroimaging studies of ASD. To better characterize our cohort, we provided a list of medications and co-occurring medical conditions (such as high blood pressure, high cholesterol, and immunological conditions) reported by all participants in Supplementary Table S5. We found that a similar number of participants in both the ASD and NT groups were on some type of prescription medication, suggesting that medications are unlikely to be the primary factor driving the group differences in BSV. Notably, 9 participants in the ASD group and 1 in the NT group reported using psychotropic medications. Although this factor could potentially influence group differences in BSV, the expected effect would likely be in the opposite direction based on findings by Lalwani et al. (2021) in light of the specific medications used in our ASD group, none of which directly target GABA. Certain medications, such as citalopram (an SSRI; Sanacora et al., 2002), amphetamines (Del Arco et al., 1999), lamotrigine, venlafaxine/effexor, and fluoxetine, have shown some evidence of increasing GABA levels (Huang et al., 2016; Mirza et al., 2005; Gören et al., 2007). Aripiprazole may increase GABA receptor expression, while risperidone tends to reduce GABA levels (Pan et al., 2016). Buspirone does not affect GABA receptors (Taylor, 1988), and the effects of amitriptyline on GABA levels remain unclear (Bang et al., 2021; Malatynska et al., 1991). In which case, the psychotropic medication use in the ASD group should make it more difficult to detect certain age-related effects. Nevertheless, we still observed a diagnosis by age interaction, with decreased BSV in the ASD group.

## Limitations

It is essential to understand age-related changes in brain function in ASD to inform best practices for cognitive and health screenings in adults with ASD and to develop interventions that may help prevent accelerated decline. Our present data relied on cross-sectional observations and a small sample size, so future work should conduct additional longitudinal analyses in larger cohorts and also investigate relationships with measures of cognitive function measures. In the future, we plan to report statistical effects related to cognition, ASD severity, and behavior measures with our findings.

## Conclusion

In our cross-sectional analyses of older adults (40-70 years) with ASD, BSV declined with age in the ASD but not NT group. Together with a previous study detecting neurotypical BSV levels in children and adolescents with ASD, our findings may indicate that decreased BSV is limited to adulthood in ASD, potentially as a result of accelerated aging. However, given limited prior research on BSV in ASD and evidence of altered GABA activity across the lifespan in ASD, additional longitudinal analyses will be necessary to determine if the results presented here truly reflect accelerated aging or arise from lifelong persistent differences in brain function.

## List of abbreviations

BSV: Bold Signal Variability

ASD: Autism Spectrum Disorder

NT: Neurotypical

RMSD: Root-Mean-Square Deviation

BMI: Body Mass Index

WASI: Wechsler Abbreviated Scale of Intelligence

VCI: Verbal Comprehension Index

PRI: Perceptual Reasoning Index

FSIQ: Full Scale Intelligence Quotient

HR: Heart Rate

RVT: Respiratory Volume per Time

TR: Repetition Time

TE: Echo Time

FOV: Field of View

PCA: Principal Component Analysis

ROI: Region of Interest

MNI: Montreal Neurological Institute

EPI: Echo Planar Imaging

CONN: Connectivity toolbox

ART: Artifact Detection Toolbox

BHFDR: Benjamini-Hochberg False Discovery Rate

SPM: Statistical Parametric Mapping.

## Declarations

### Ethics approval and consent to participate Availability of data and materials

Data included in these analyses will be available through the National Institute of Mental Health Data Archive (nda.nih.gov/). Researchers who qualify may request access through this system.

## Acknowledgements

Our sincere thanks to our participants and their families for sharing their time with us.

## Footnotes

Throughout this document, we used terms such as “adults on the autism spectrum” and “neurotypical” (NT) peers and acknowledge that such language may not be preferred by everyone. We selected terminology that we believed would be more widely used by the individuals represented in this study, aiming to move towards identity-first language within the context of a neurodiversity framework.

