## Supplementary Materials for "Decreased BOLD signal variability in middle-aged and older adults on the Autism Spectrum"

**Supplementary Material:**

**Table S1. Main effect of Age** (all tested ROIs shown) - (selected based on findings reported by Lalwani et al. 2021)

| **ROIs** | **t-value** | **p-value** | **BFDR** |
| --- | --- | --- | --- |
| **Frontal Orbital Cortex (L)** | -1.187 | 0.240 | 0.280 |
| **Frontal Orbital Cortex (R)**** | -3.361 | 0.001 | 0.017 |
| **Frontal Pole (L)** | -1.465 | 0.148 | 0.213 |
| **Frontal Pole (R)** | -1.337 | 0.186 | 0.240 |
| **Inferior Frontal Gyrus, pars opercularis (L)** | -1.614 | 0.112 | 0.168 |
| **Inferior Frontal Gyrus, pars opercularis (R)**** | -2.822 | 0.006 | 0.046 |
| **Inferior Frontal Gyrus, pars triangularis (L)*** | -2.552 | 0.013 | 0.079 |
| **Inferior Frontal Gyrus, pars triangularis (R)** | -1.769 | 0.082 | 0.155 |
| **Paracingulate Gyrus (L)** | -1.347 | 0.183 | 0.240 |
| **Paracingulate Gyrus (R)** | -1.975 | 0.053 | 0.125 |
| **Insular Cortex (L)*** | -2.494 | 0.015 | 0.079 |
| **Insular Cortex (R)**** | -3.965 | <0.001 | 0.007 |
| **Precentral Gyrus (L)** | -1.613 | 0.112 | 0.167 |
| **Precentral Gyrus (R)** | -1.164 | 0.249 | 0.280 |
| **Superior Frontal Gyrus (L)*** | -2.070 | 0.043 | 0.125 |
| **Superior Frontal Gyrus (R)*** | -2.294 | 0.025 | 0.101 |
| **Superior Parietal Lobule (L)** | -1.177 | 0.243 | 0.280 |
| **Superior Parietal Lobule (R)** | -1.951 | 0.056 | 0.125 |
| **Temporal Pole (L)** | -1.695 | 0.095 | 0.163 |
| **Temporal Pole (R)** | -1.335 | 0.187 | 0.240 |
| **Planum Temporale (L)** | -1.295 | 0.200 | 0.248 |
| **Planum Temporale (R)** | -0.656 | 0.514 | 0.561 |
| **Inferior Temporal Gyrus, anterior division (L)** | -1.823 | 0.073 | 0.146 |
| **Inferior Temporal Gyrus, anterior division (R)** | -0.295 | 0.769 | 0.769 |
| **Temporal Occipital Fusiform Cortex (L)**** | -3.332 | 0.001 | 0.017 |
| **Temporal Occipital Fusiform Cortex (R)*** | -2.112 | 0.039 | 0.125 |
| **Inferior Temporal Gyrus, temporooccipital part (L)*** | -2.219 | 0.030 | 0.108 |
| **Inferior Temporal Gyrus, temporooccipital part (R)*** | -2.389 | 0.020 | 0.089 |
| **Inferior Temporal Gyrus, posterior division (L)** | -1.715 | 0.091 | 0.163 |
| **Inferior Temporal Gyrus, posterior division (R)** | -0.410 | 0.683 | 0.703 |
| **Lateral Occipital Cortex, superior division (L)** | -1.617 | 0.111 | 0.167 |
| **Lateral Occipital Cortex, superior division (R)** | -1.962 | 0.054 | 0.125 |
| **Lateral Occipital Cortex, inferior division (L)** | -1.893 | 0.063 | 0.133 |
| **Lateral Occipital Cortex, inferior division (R)**** | -2.834 | 0.006 | 0.046 |
| **Cuneal Cortex (L)** | -0.437 | 0.664 | 0.703 |
| **Cuneal Cortex (R)*** | -2.018 | 0.048 | 0.125 |
| Note: * denote p-values <0.05, **p-values remain significant after as Benjamini-Hochberg False Discovery Rate (BFDR) adjustment. | | | |

**Table S2. Main effects of Diagnosis** (all tested ROIs shown)

| **ROIs** | **t-value** | **p-value** | **BFDR** |
| --- | --- | --- | --- |
| **Frontal Orbital Cortex (L)** | -1.079 | 0.285 | 0.351 |
| **Frontal Orbital Cortex (R)** ** | -3.219 | 0.002 | 0.024 |
| **Frontal Pole (L)** | -1.508 | 0.137 | 0.223 |
| **Frontal Pole (R)** | -1.349 | 0.182 | 0.262 |
| **Inferior Frontal Gyrus, pars opercularis (L)** | -1.299 | 0.199 | 0.275 |
| **Inferior Frontal Gyrus, pars opercularis (R)** * | -2.471 | 0.016 | 0.101 |
| **Inferior Frontal Gyrus, pars triangularis (L)** | -2.411 | 0.019 | 0.101 |
| **Inferior Frontal Gyrus, pars triangularis (R)** | -1.570 | 0.121 | 0.220 |
| **Paracingulate Gyrus (L)** | -0.966 | 0.338 | 0.392 |
| **Paracingulate Gyrus (R)** | -1.390 | 0.170 | 0.254 |
| **Insular Cortex (L)** | -2.321 | 0.024 | 0.105 |
| **Insular Cortex (R)** ** | -3.660 | 0.001 | 0.016 |
| **Precentral Gyrus (L)** | -1.568 | 0.122 | 0.220 |
| **Precentral Gyrus (R)** | -1.061 | 0.293 | 0.351 |
| **Superior Frontal Gyrus (L)** | -1.771 | 0.081 | 0.172 |
| **Superior Frontal Gyrus (R)** | -2.024 | 0.047 | 0.124 |
| **Superior Parietal Lobule (L)** | -1.093 | 0.278 | 0.351 |
| **Superior Parietal Lobule (R)** * | -2.015 | 0.048 | 0.124 |
| **Temporal Pole (L)** | -1.539 | 0.129 | 0.221 |
| **Temporal Pole (R)** | -1.183 | 0.241 | 0.321 |
| **Planum Temporale (L)** | -0.792 | 0.431 | 0.485 |
| **Planum Temporale (R)** | -0.250 | 0.803 | 0.826 |
| **Inferior Temporal Gyrus, anterior division (L)** | -1.716 | 0.091 | 0.182 |
| **Inferior Temporal Gyrus, anterior division (R)** | -0.067 | 0.947 | 0.947 |
| **Inferior Temporal Gyrus, posterior division (L)** | -1.776 | 0.081 | 0.172 |
| **Inferior Temporal Gyrus, posterior division (R)** | -0.415 | 0.679 | 0.741 |
| **Temporal Occipital Fusiform Cortex (L)** ** | -3.485 | 0.001 | 0.016 |
| **Temporal Occipital Fusiform Cortex (R)** * | -2.168 | 0.034 | 0.122 |
| **Inferior Temporal Gyrus, temporooccipital part (L)** | -2.274 | 0.026 | 0.105 |
| **Inferior Temporal Gyrus, temporooccipital part (R)** | -2.394 | 0.020 | 0.101 |
| **Lateral Occipital Cortex, superior division (L)** | -1.475 | 0.145 | 0.227 |
| **Lateral Occipital Cortex, superior division (R)** * | -2.038 | 0.046 | 0.124 |
| **Lateral Occipital Cortex, inferior division (L)** * | -2.079 | 0.042 | 0.124 |
| **Lateral Occipital Cortex, inferior division (R)** ** | -2.941 | 0.005 | 0.041 |
| **Cuneal Cortex (L)** | -0.385 | 0.701 | 0.743 |
| **Cuneal Cortex (R)** * | -1.960 | 0.054 | 0.131 |
| Note: * denote p-values <0.05, **p-values remain significant after Benjamini-Hochberg False Discovery Rate (BFDR) adjustment. | | | |

**Table S3. Age-by-Diagnosis** (all tested ROIs shown)

| **ROIs** | **t-value** | **p-value** | **BFDR** |
| --- | --- | --- | --- |
| Frontal Orbital Cortex (L) | 0.532 | 0.597 | 0.643 |
| Frontal Orbital Cortex (R)** | 3.095 | 0.003 | 0.031 |
| Frontal Pole (L) | 1.270 | 0.209 | 0.286 |
| Frontal Pole (R) | 1.191 | 0.238 | 0.298 |
| Inferior Frontal Gyrus, pars opercularis (L) | 1.328 | 0.189 | 0.284 |
| Inferior Frontal Gyrus, pars opercularis (R)* | 2.473 | 0.016 | 0.100 |
| Inferior Frontal Gyrus, pars triangularis (L)* | 2.234 | 0.029 | 0.100 |
| Inferior Frontal Gyrus, pars triangularis (R) | 1.647 | 0.105 | 0.209 |
| Paracingulate Gyrus (L) | 1.010 | 0.316 | 0.356 |
| Paracingulate Gyrus (R) | 1.519 | 0.134 | 0.220 |
| Insular Cortex (L)* | 2.369 | 0.021 | 0.100 |
| Insular Cortex (R)** | 3.764 | 0.000 | 0.013 |
| Precentral Gyrus (L) | 1.729 | 0.089 | 0.192 |
| Precentral Gyrus (R) | 1.254 | 0.214 | 0.286 |
| Superior Frontal Gyrus (L) | 1.719 | 0.090 | 0.192 |
| Superior Frontal Gyrus (R)* | 2.023 | 0.047 | 0.131 |
| Superior Parietal Lobule (L) | 1.186 | 0.240 | 0.298 |
| Superior Parietal Lobule (R)* | 1.958 | 0.055 | 0.131 |
| Temporal Pole (L) | 1.581 | 0.119 | 0.220 |
| Temporal Pole (R) | 1.281 | 0.205 | 0.286 |
| Planum Temporale (L) | 1.028 | 0.308 | 0.356 |
| Planum Temporale (R) | 0.517 | 0.607 | 0.643 |
| Inferior Temporal Gyrus, anterior division (L) | 1.516 | 0.134 | 0.220 |
| Inferior Temporal Gyrus, anterior division (R) | 0.094 | 0.925 | 0.925 |
| Inferior Temporal Gyrus, posterior division (L) | 1.539 | 0.129 | 0.220 |
| Inferior Temporal Gyrus, posterior division (R) | 0.432 | 0.668 | 0.687 |
| Temporal Occipital Fusiform Cortex (L)** | 3.467 | 0.001 | 0.017 |
| Temporal Occipital Fusiform Cortex (R)* | 2.199 | 0.032 | 0.100 |
| Inferior Temporal Gyrus, temporooccipital part (L)* | 2.241 | 0.029 | 0.100 |
| Inferior Temporal Gyrus, temporooccipital part (R)* | 2.337 | 0.023 | 0.100 |
| Lateral Occipital Cortex, superior division (L) | 1.484 | 0.143 | 0.224 |
| Lateral Occipital Cortex, superior division (R)* | 1.987 | 0.051 | 0.131 |
| Lateral Occipital Cortex, inferior division (L)* | 2.206 | 0.031 | 0.100 |
| Lateral Occipital Cortex, inferior division (R)** | 3.043 | 0.003 | 0.031 |
| Cuneal Cortex (L) | 0.532 | 0.597 | 0.643 |
| Cuneal Cortex (R)* | 2.177 | 0.033 | 0.100 |
| *Note:* * denote p-values <0.05, **p-values remain significant after Benjamini-Hochberg False Discovery Rate (BFDR) adjustment. | | | |

**Table S4.** Health Measures - Frequencies

|  | **NT** | **ASD** | **Statistics** |
| --- | --- | --- | --- |
| High Blood Pressure (no/yes) | 23/4 | 13/6 | x^2^= 1.8, *p*=0.2 |
| High Cholesterol (no/yes) | 22/5 | 11/8 | x^2^= 3.7, *p*=0.08 |

*Data were missing for several participants:* HBP and HC (12 NT and 9 ASD).

**Table S5.** Medication List for All Participants

| ID | Dx | HBP | HC | Medications |
| --- | --- | --- | --- | --- |
| 1 | NT | Yes | Yes | None |
| 2 | NT | No | Yes | None |
| 3 | NT | No | No | ActivatedYou (daily supplement); Turmeric (probiotic); Vitamin D 3000 IU (inflammation) |
| 4 | NT | No | No | Aspirin (as needed, OTC); Naproxen (as needed, OTC); Zyrtec (as needed, OTC) |
| 5 | NT | Yes | No | Aspirin (as needed); Benadryl (as needed) |
| 6 | NT | No | Yes | Citalopram 20mg (daily, depression); Rosuvastatin 10mg (daily, HC); Tamsulosin 8mg (daily, urinary retention); Omeprazole 20mg (daily, stomach issues); Naproxen 500mg (as needed, anti-inflammatory) |
| 7 | NT | Yes | Yes | Flomax 0.8mg (2 capsules, daily, urinary retention); Prilosec 20mg (daily, proton-pump inhibitor); Zyrtec 10mg (daily, allergy); Prinivil 20mg (daily, ACE inhibitor); Travatan (eye drops, daily, glaucoma); Lipitor 40mg (daily, HC); Ultram 50mg (every 6-8 hours, pain); Finasteride 5mg (daily, urinary retention) |
| 8 | NT | No | No | Ibuprofen (as needed) |
| 9 | NT | No | No | Ibuprofen (as needed); Claritin/Benadryl (OTC, allergy) |
| 10 | NT | No | No | Levothyroxine 125mg (daily, thyroid replacement); Ibuprofen 400mg (2x/month, headache) |
| 11 | NT | Yes | No | Lisinopril (daily, HBP) |
| 12 | NT | No | No | Mucinex DM (2x daily, mucus); Cetirizine Hydrochloride 10mg (OTC, cough) |
| 13 | NT | No | Yes | Simvastatin 10mg (daily, HC); Aspirin 81mg (daily, heart health) |
| 14 | NT | No | No | Vitamin D (OTC); Vitamin C |
| 15 | ASD | Yes | Yes | Adderall XR 60mg (daily, focus, calming); Effexor 220mg (daily, depression); Testosterone 200mg (weekly, low testosterone); Anastrozole 1mg (weekly, estrogen blocker); Levothyroxine 100mcg (daily, hypothyroidism); Chlorthalidone 25mg (daily, diuretic); Tramadol 50mg (sleep) |
| 16 | ASD | No | Yes | Atorvastatin (HC); Benztropine (anti-tremor); Risperidone (irritability) |
| 17 | ASD | No | Yes | Buspirone 15mg (2x, anxiety); Simvastatin 20mg (1x, HC); Trazodone 300mg (1x, sleep); Effexor 150mg (1x, anxiety); Lyrica 75mg (2x daily, nerve pain); Methocarbamol 750mg (3x, muscle relaxant) |
| 18 | ASD | No | No | Calcium 600mg (1x daily, bone health); Vitamin D3 800 IU (daily) |
| 19 | ASD | Yes | Yes | Enalapril (BP, once daily); Simvastatin (HC, as prescribed) |
| 20 | ASD | Yes | No | Hydrochlorothiazide 25mg (once daily, HBP); Lisinopril 40mg (once daily, HBP) |
| 21 | ASD | No | Yes | Lamotrigine 50mg (daily, depression); Abilify 5mg (daily, mania) |
| 22 | ASD | No | No | Loratadine (as needed, allergies); Albuterol (as needed, asthma) |
| 23 | ASD | Yes | No | Losartan 100mg (HBP) |
| 24 | ASD | No | Yes | Pravastatin 20mg (daily, HC) |
| 25 | ASD | Yes | No | Prozac (anxiety) |
| 26 | ASD | No | Yes | Synthroid 175mcg (daily, hypothyroidism); Amitriptyline 50mg (daily, chronic pain); Metoprolol Tartrate 100mg (2x daily, BP); Aspirin 81mg (daily, blood thinner); Rosuvastatin 40mg (daily, HC) |
| 27 | ASD | No | No | Tamoxifen (daily, cancer therapy); Gabapentin (daily, restless leg syndrome); Effexor (daily, depression) |
| 28 | ASD | Yes | Yes | Tamsulosin (HBP, HC); Atorvastatin (HBP, HC); Lisinopril (HBP, HC); Flonase (sinus issues) |
| 29 | ASD | No | No | Valacyclovir 500mg (2x per day, recurrent ocular herpes); Loratadine 10mg (1x daily, allergies); Breo Ellipta (200mcg/25mcg, 1x daily, asthma); Albuterol Sulfate (90mcg, as needed, asthma); Adderall 30mg (2x daily, ADHD); Previously on anxiolytics, antidepressants, mood stabilizers |
| 30 | ASD | Yes | No | Valsartan (daily, BP); Zyrtec (allergies); Sudafed (sinuses); Fluticasone (as needed, sinusitis); Azelastine (as needed, sinusitis); Methylphenidate (ADHD); Lamotrigine (as needed, anxiety/hypomania) |
| 31 | ASD | No | No | Vesicare (bladder relaxant); Prevacid (proton-pump inhibitor); OTC allergy medication |
| *Note:* High Blood Pressure=HBP, High Cholesterol=HC, Immunological Conditions=IMM, Over The Counter=OTC. All medicines are prescribed unless otherwise stated. | | | | |
